# Primary metabolism underpins the execution of immune responses in different tissues of the same plant

**DOI:** 10.1101/2025.10.11.681807

**Authors:** Emma C. Raven, Rhea Stringer, Catherine Walker, Hannah Rae Thomas, Christine Faulkner

## Abstract

Cell-surface perception of microbes triggers a range of rapid immune responses in plants that include the induction of cell isolation by plasmodesmal closure, the production of reactive oxygen species, and changes to gene expression. Here, we identify that some of these immune response are differentially executed in leaves of different ages in the same plant. We observed that when compared to mature leaves, young, expanding leaves do not close their plasmodesmata, have a reduced transcriptional response to immune elicitors, and are more susceptible to a bacterial pathogen. Disconnecting leaf age from physiology, we determined that both plasmodesmal closure and the magnitude of transcriptional responses are dependent on whether the leaf is a carbon sink or a carbon source. To probe the relevance of differential regulation of plasmodesmata in sink and source tissues, we forced plasmodesmal closure in young sink leaves and found this perturbed the normal outputs of growth and defence. Thus, we propose that sink leaves do not close their plasmodesmata during immune reponses to prioritise carbon use for growth over defence, and consequently that primary metabolism underpins different immune response profiles in different leaves of the same plant.

## Introduction

The plant immune system relies on a suite of receptors to perceive microbial molecules intracellularly and at the cell surface. Cell surface receptors (pattern recognition receptors, PRRs) are anchored to the plasma membrane where they bind to extracellular microbial molecules collectively called pattern associated molecular patterns (PAMPs). Thus, binding between receptors and their ligands triggers immune responses that include changes to gene expression, the production of reactive oxygen species (ROS), a calcium ion influx, and closure of plasmodesmata, which are the intercellular cytoplasmic connections between cells. These responses collectively enhance cellular defence against microbes and in some cases offer effective resistance^1^.

PRRs activate diverging signalling cascades that ultimately lead to execution of the different immune responses. For example, PRRs transmit signals via cytoplasmic kinases to the plasma membrane-located NADPH oxidase RESPIRATORY BURST OXIDASE HOMOLOGUE D (RBOHD) to produce a ROS burst and, via mitogen-activated protein kinases (MAPKs), induce changes in gene expression^1^. PRRs also activate the independent production of the β-1,3-glucan callose via CALLOSE SYNTHASES (CalSs) at plasmodesmata to mediate plasmodesmal closure^2-4^ and in macroscopic deposits across a tissue^5^. In plasmodesmata, CalSs are activated by specific signalling cascades that utilise plasmodesmata-specific machinery^6-8^. The independence of immune signalling at plasmodesmata from other PAMP-triggered responses suggests the plasmodesmal response can be differentially regulated, raising questions regarding when this might occur and how it is beneficial to overall physiological success.

Plasmodesmata enable the intercellular exchange of soluble molecules such as sugars, proteins, and nucleic acids between cells, tissues and organs meaning their dynamic aperture can regulate molecular exchange across spatiotemporal scales. Mutants with impaired PAMP-triggered plasmodesmal closure are more susceptible to a range of pathogens^2,9,10^, suggesting that immune responses are enhanced by cell isolation. Additionally, diverse pathogens target plasmodesmal signalling pathways to enhance infection^11,12^, possibly by maintaining cell-to-cell connectivity to enable the transmission of pathogen molecules through the host and the delivery of resources to the infection site. Therefore, plasmodesmal connectivity between cells is critical for both steady-state plant physiology as well as responses to pathogens and other environmental changes.

Across a growing plant, there are distinct organs and tissues at different stages of development. The heterogeneity in developmental age across an individual has been previously identified to interact with stress responses, with leaves of different ages exhibiting different activities of the salicylic acid activated transcription factor PBS3 to optimise reproductive success in the presence of stresses^13^. Carbon physiology also varies across an individual plant, e.g. expanding leaves are carbon sinks while mature expanded leaves are carbon sources. Carbon sinks unload sugars from sieve elements into surrounding cells through open plasmodesmata^8^, fuelling respiration to drive growth and development^9,10^. As sink leaves develop, rates of carbon assimilation increase until the net carbon gain exceeds the loss of carbon through respiration, at which point leaves transition to become carbon source tissues^10-12^.

To explore how different developmental and physiological states impact immunity, we assessed the regulation of immune responses in leaves of Arabidopsis of different ages and carbon status. Examining cell surface activated immunity, we found that while the PAMP-triggered ROS burst and macroscopic callose deposition were executed normally in young sink leaves, these leaves do not close their plasmodesmata in response to stress, are more susceptible to a bacterial pathogen and show a reduced transcriptional immune response relative to mature expanded leaves. However, inducing the sink-to-source transition enables plasmodesmal responses to elicitors and enhances transcriptional activity in young leaves, indicating that carbon physiology regulates these elements of immunity. As phloem unloading occurs via plasmodesmata, our data suggests independent regulation of plasmodesmal immune responses allows tissues to balance the demand for carbon in growth against the benefits of cell isolation during defence.

## Results

### Young leaves can perceive and respond to PAMPs, but show a reduced transcriptional response

To explore whether PAMP-triggered immune responses vary throughout developmental and physiological stages, we investigated a subset of immune responses in expanded, mature leaves and recently emerged, expanding young leaves of 5-week-old *Arabidopsis thaliana* (Arabidopsis) plants (Fig. 1A). We first assayed for the PAMP-triggered, apoplastic reactive oxygen species (ROS) burst using a luminol-based assay. Both chitin and flg22 triggered a rapid ROS burst in young and mature leaves (Fig. 1B and C, Fig. S1A, B). While young leaves produced higher cumulative relative light units (RLU), a measure of the overall reactivity of the system, it is unclear whether this relates to the magnitude of the response or technical differences in the uptake of reagents in young leaves. We also measured macroscopic callose deposition in response to flg22. Using live-imaging of aniline blue stained leaves, we detected flg22-triggered macroscopic callose deposition in both young and mature leaves (Fig. 1C, D), indicating young leaves can synthesise and accumulate callose in response to elicitors. Overall, these results indicate that young leaves can perceive and respond to PAMPs.

**Figure 1:**
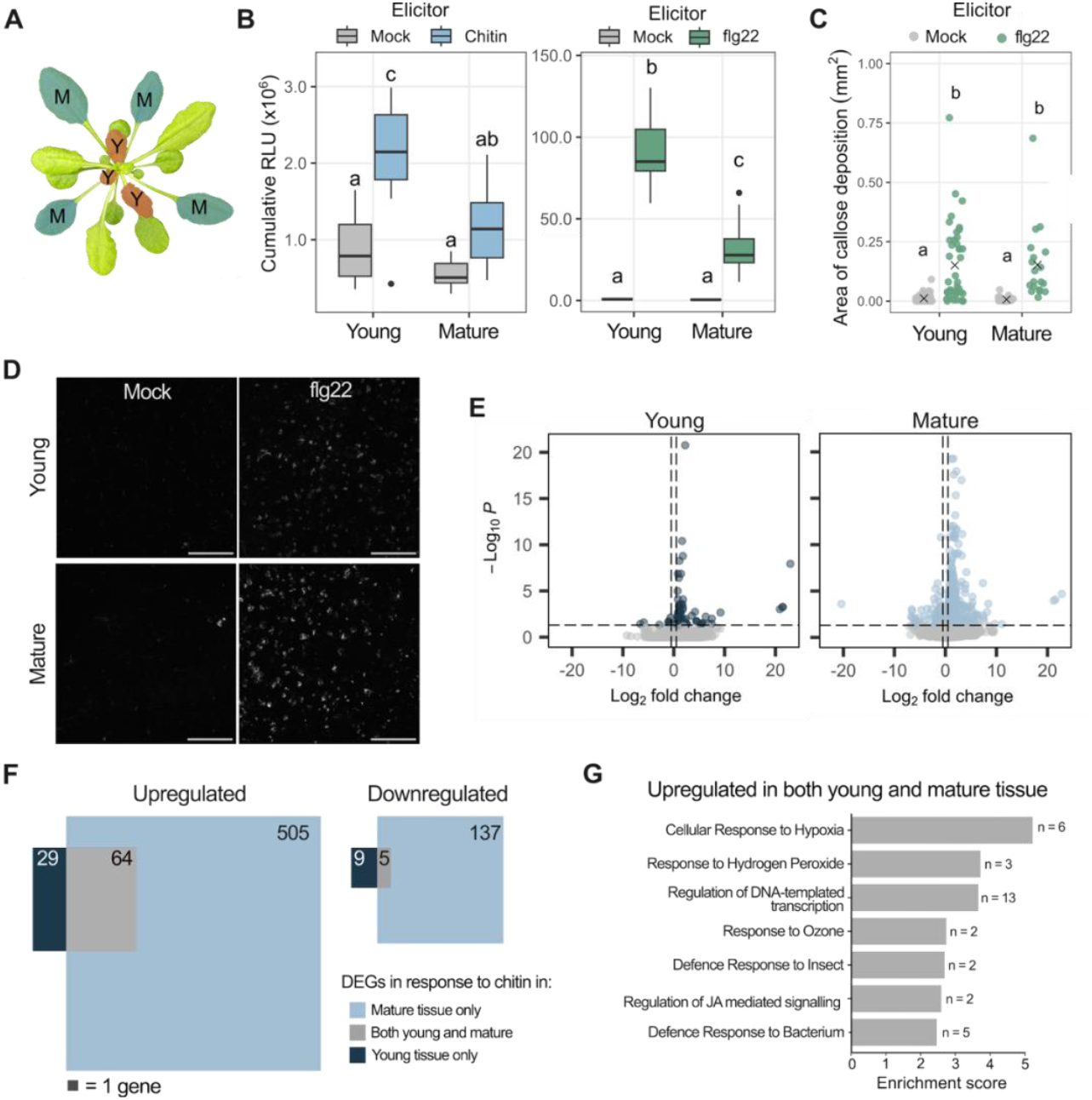
Young leaves can perceive and respond to PAMPs. (A) Mature (M) and Young (Y) leaves sampled on 5-week-old Arabidopsis rosettes. (B) Cumulative Relative Light units (RLU) produced from young and mature leaves in response to chitin (blue) and flg22 (green) treatment compared to mock (grey). Different letters indicate p < 0.05, n = 12. (C) Area per image covered by aniline blue stained callose in young and mature leaves in response to mock (grey) or flg22 (blue) treatment. ‘X’ indicates the median. Different letters indicate p < 0.05, n = 16. (D) Representative confocal images of aniline blue stained macroscopic callose deposition in young and mature tissue in response to mock or flg22 treatment. Scale bars are 200 µm (E) Differentially expressed genes (DEGs, p_adj_<0.05, log_2_[fold change] > |0.5|) in response to chitin compared to water treatment. (F) Venn diagram of chitin-triggered DEGs in mature tissue (light blue), young tissue (dark blue) and both (grey). (G) Enriched Gene Ontology (GO) terms for genes upregulated in both mature and young tissue, n indicates the number of genes.

To further explore PAMP-triggered immune responses in young leaves, we analysed chitin-induced transcriptional responses in young and mature leaf tissue. We used RNA-sequencing (seq) to profile differentially expressed genes (DEGs) 30 minutes post-chitin treatment and found that young leaves have a much smaller transcriptional response to chitin compared to mature leaf tissue. While mature leaves showed differential expression of 712 genes in response to chitin, young leaves only differentially expressed 103 genes (Fig. 1E, Supplemental Dataset S1). Despite young leaves having fewer DEGs, there was a set of 64 up-regulated DEGs common to young and mature leaf tissue (Fig. 1F, Supplemental Dataset S1). A Gene Ontology (GO)-term enrichment analysis of these genes identified GO terms associated with defense against insects and bacteria as well as jasmonic acid signalling, suggesting immune responses occur in both young and mature leaves (Fig. 1G, Supplemental Dataset S1). This suggests young leaves transcriptionally respond to chitin, albeit to a lesser degree than mature leaves.

### Young leaves do not close their plasmodesmata in response to PAMPs

To date, our research into PAMP-triggered plasmodesmal closure has been focused on the response in mature, expanded leaves of 5-week-old Arabidopsis^2-4^. To determine if this response oocurs in younger leaves, we assayed for PAMP-induced reductions in the cell-to-cell movement of GFP in expanding young leaves using microprojectile bombardment assays (Fig. 2A, Supplemental Dataset S2). Consistent with previous studies^2,10^, chitin and flg22 reduced the spread of GFP from bombarded cells in mature leaves indicating plasmodesmal closure (Fig. 2B-C, Supplemental Dataset S2). However, young leaves did not exhibit chitin- or flg22-triggered plasmodesmal closure. To determine whether this lack of response was specific to PAMPs, we tested other stress elicitors, and observed that while salicylic acid (SA), hydrogen peroxide (H_2_O_2_) and abscisic acid (ABA) induced plasmodesmal closure in mature leaves^14,15^, they did not in young leaves (Fig. 2D, E, Supplemental Dataset S2). Collectively, this suggests that plasmodesmata in young leaves do not respond to stress.

**Figure 2:**
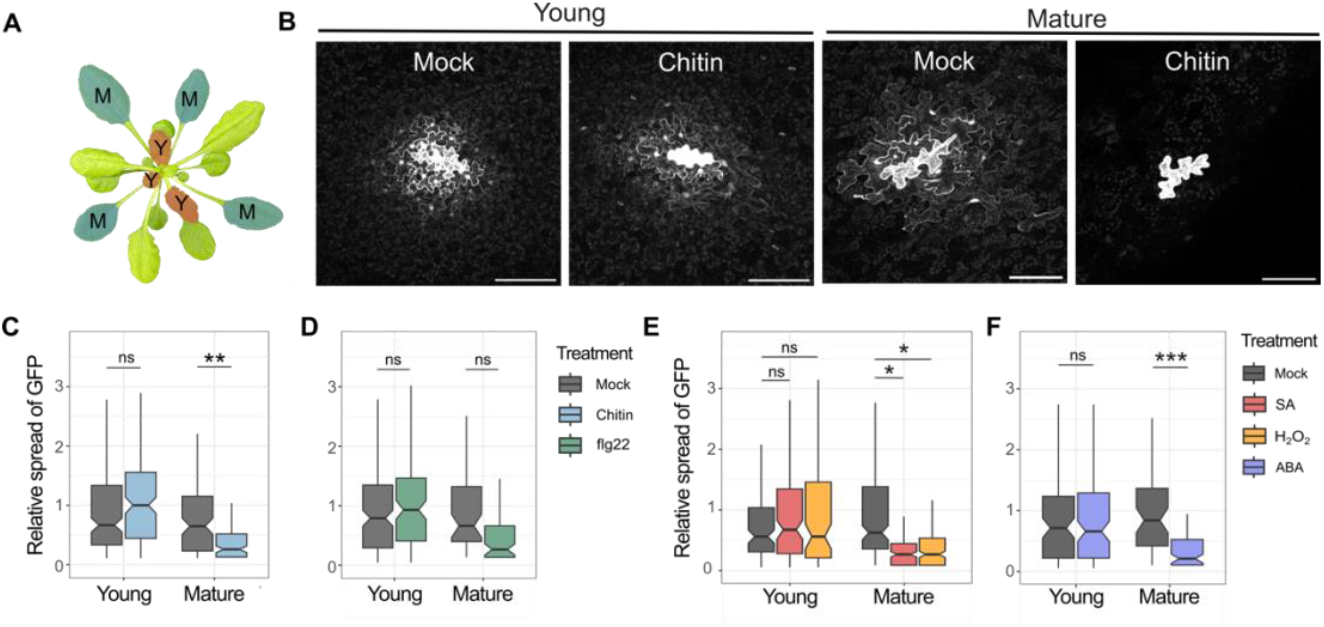
Plasmodesmal closure does not occur in young leaves. (A) Representative micrographs showing GFP fluorescence from single cell transformation events in mock and chitin treated young and mature leaves, scale bars indicate 100 µm. (B-G) Relative spread of GFP in leaves treated with water (mock, grey) and (B) chitin (light blue), n > 88, (C) flg22 (green), n > 109, (D) salicylic acid (SA, red) and hydrogen peroxide (H_2_O_2_, orange), n > 93, and (E) abscisic acid (ABA, purple) n > 53. Asterisks indicate statistical significance compared to the mock treatment: *p < 0.05, **p < 0.01, ***p < 0.001.

Plasmodesmal responses to PAMPs are mediated by specific signalling machinery^2-4,10^ and it is possible that young leaves do not close their plasmodesmata because they do not produce these proteins. Therefore, we mined transcriptional data from young leaves to identify plasmodesmal signalling components. Chitin-triggered plasmodesmal closure relies on *LYM2, RBOHD, NHL3, PDLP1* and *PDLP5*^2-4^, and each of these genes was similarly expressed in both young and mature leaves (Fig. S1C). Additionally, LYM2 localises to the plasmodesmata in young leaves when *LYM2-citrine* is expressed from the *LYM2* native promoter (Fig. S1D). Therefore, we conclude that young leaves have the machinery to close plasmodesmata but that the signalling pathway is either not activated or efficiently executed.

### Covering mature leaves induces a carbon sink-to-source transition in young leaves

It is likely that mature expanded leaves are carbon source tissue while young, expanding leaves are carbon sinks. To address whether the sink or source status underpins plasmodesmal responsiveness, we developed a simple experimental system to decouple developmental age from the direction of net carbon transport. For this, we covered the mature leaves of an Arabidopsis rosette with foil (Fig. 3A), limiting the light reactions of photosynthesis to reduce the assimilation and export of photosynthates. We predicted the uncovered young sink leaves would receive little or no import of carbon from covered mature leaves and so would transition to become developmentally young source tissue.

**Figure 3:**
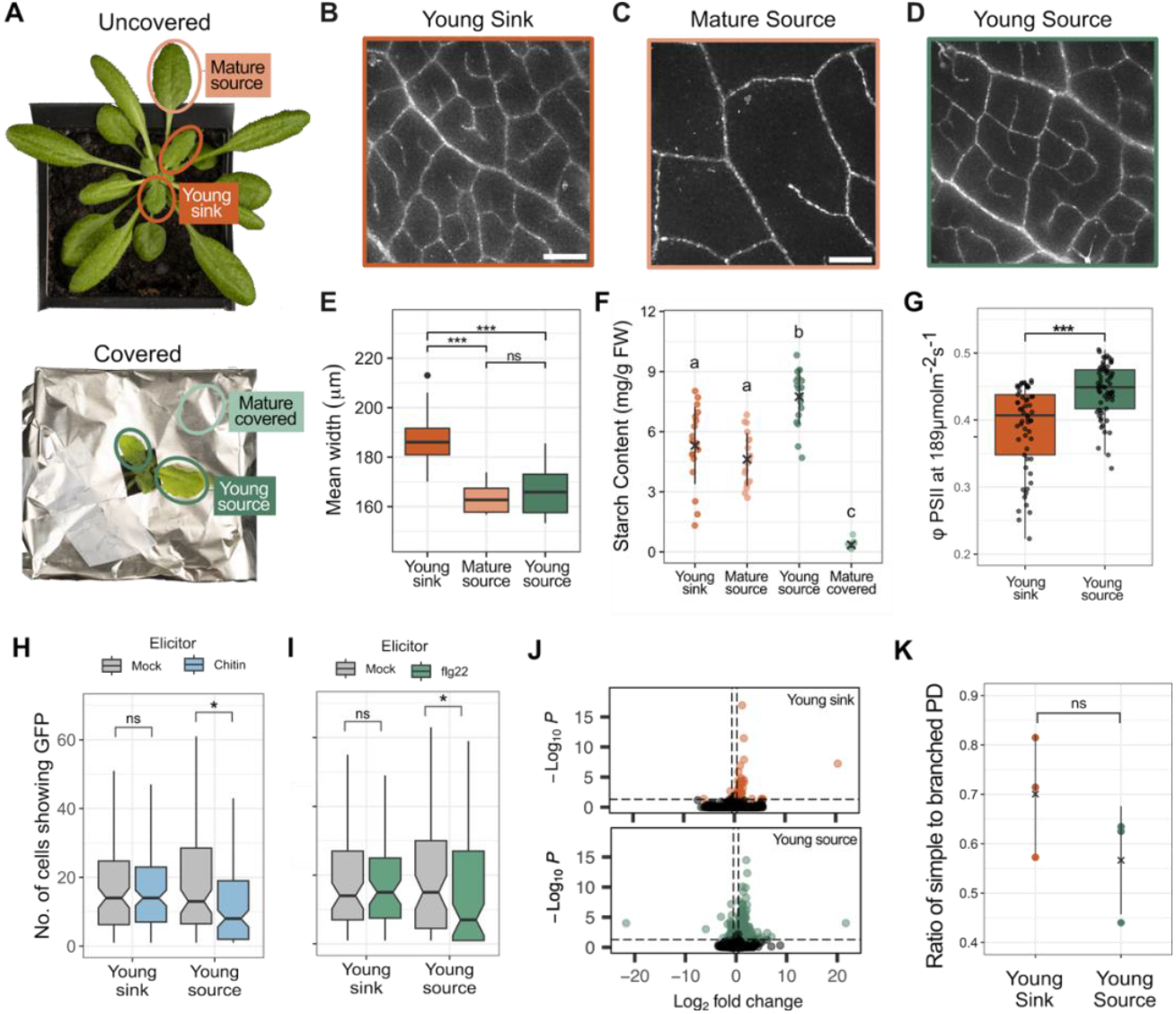
Inducing the sink-to-source transition changes plasmodesmal responsiveness. (**A**) 5 week old Arabidopsis rosettes for which the mature leaves (top) have been covered to induce the sink-to-source transition in young leaves (bottom). (**B**) Extended depth of focus images of GFP movement in young sink, (**C**) mature source, and (**D**) young source leaves of pAtSUC2::GFP plants, scale bars indicate 500 µm. (**E**) Quantification of the distance moved by GFP out of the phloem 24 hours post covering, n > 13. (**F**) Quantification of the starch content in leaves 24 hours following covering mature leaves, n = 18. (**G**) Quantification of the operating efficiency of photosystem II (φPSII) in leaves, n > 12. (**H**) Movement of GFP from single cells in microprojectile bombardment assays in young sink and source leaves treated with chitin (blue, n > 105) and (**I**) flg22 (green, n > 64). (**J**) Volcano plots of differentially expressed genes in response to chitin compared to water treatment in young sink and young source tissue, where orange and green points indicate padj<0.05, log2(fold change) > |0.5|. (**K**) Ratio of simple to branched plasmodesmata (PD) in young sink and young source leaves. Asterisks indicate statistical significance compared to the mock treatment: *p < 0.05, **p < 0.01, ***p < 0.001, and no-significance in ‘ns’. Different letters indicate p < 0.05.

To verify that covering mature leaves induced a physiological transition in young leaves, we assayed for reduced phloem unloading in young leaves of covered plants. For this we used *pAtSUC2::GFP* lines that produce GFP in phloem companion cells^16^. In these plants, GFP fluorescence spreads from the veins as the GFP unloads from the phloem into surrounding cells^17,18^. To quantify this, we imaged leaves 24 and 48 hours after covering mature leaves and measured the distance GFP fluorescence is detected from the vein^19^. This distance was reduced in young leaves from covered plants relative to young leaves from uncovered plants at both timepoints (Fig. 3B-E, S2A), indicating that phloem unloading has reduced or ceased and that young leaves from covered plants show characteristics of source tissues (hereafter ‘young source leaves’).

To further profile the changes in young source leaves, we quantified their starch and soluble sugar contents. 24 and 48 hours after covering, young source leaves showed a significantly higher accumulation of starch at the end of day than in young sink and mature source leaves (Fig. 3F, S2C). No changes in soluble sugars were observed in young leaves 24 hours after covering (Fig. S2B), but glucose content decreased in young source leaves 48 hours after covering relative to young sink leaves (Fig. S2C). Covered mature leaves had very low levels of starch and soluble sugars indicating their storage pools were depleted and that they were unlikely to supply sugars to young leaves.

We measured pulsed amplitude modulation (PAM) chlorophyll fluorescence to determine the operating efficiency of Photosystem II (ΦPSII) in photosynthetic active radiation (PAR) in different leaves. Young source leaves showed a significant increase in ΦPSII at ambient light conditions (189 µmol m^−2^ s^−1^) compared to young sink leaves (Fig. 3G). Although in C_3_ plants it is not possible to directly relate ΦPSII to carbon assimilation without further characterisation of the electron transport chain^20^, the changes to ΦPSII, alongside the changes in starch accumulation and the cessation of GFP unloading from the phloem, indicate physiological changes have occurred in young leaves of covered plants consistent with increased carbon assimilation. Overall, these physiological changes suggest covering the mature leaves forces the young leaves to transition from carbon sink to source physiology.

### Young source leaves exhibit PAMP-triggered plasmodesmal closure and enhanced transcriptional responses

To determine whether the sink or source status of a leaf underpins the observed difference in PAMP-triggered plasmodesmal responses, we assayed plasmodesmal responses in young source leaves. Thus, we performed microprojectile bombardment assays on young leaves 24 hours post-covering of the mature leaves of the rosette. In contrast to young sink leaves, young source leaves showed a reduction in the cell-to-cell mobility of GFP when treated with chitin and flg22 (Fig. 3H, Supplemental Dataset S2), indicating that young leaves can close their plasmodesmata in response to PAMPs if they transition to carbon source status.

Additionally, we assayed for a transcriptional response to chitin in young source and young sink leaves, profiling DEGs 30 minutes post-chitin treatment. 24 hours after covering mature leaves, RNA-seq revealed that young source leaf tissue has a greater transcriptional response to chitin compared to young sink leaf tissue. Young source leaves differentially expressed 337 genes in response to chitin, while young sink leaves differentially expressed only 62 genes (Fig. 3J, Supplemental Dataset S1), with 52 genes commonly differentially expressed (Fig. S2F, Supplemental Dataset S1). A GO-term enrichment analysis on upregulated genes in young sink and source leaves showed defence-related GO-terms were enriched in both tissue types (Fig. S2D, E). This suggests that the chitin response in young source leaves differs to young sink lives in magnitude but not in nature.

### Plasmodesmal responses are not associated with plasmodesmal structure

Plasmodesmal structure increases in complexity during the sink-to-source transition^21,22^, i.e. sink leaves have predominantly simple plasmodesmata while source leaves have predominantly branched plasmodesmata. To explore whether plasmodesmal structure correlates with stress responsiveness, we aimed to profile plasmodesmal closure in lines in which a *Potato Leaf Roll Virus* movement protein 17 (MP17)-GFP fusion marks branched plasmodesmata. We reasoned we could track cells with and without branched plasmodesmata using this marker^21,23^, but found constitutive expression of *MP17-GFP* significantly reduced cell-to-cell movement of GFP in mature leaves (Fig. S3A-C, Supplemental Dataset S2), making the assay impossible.

As an alternative approach, we asked whether simple plasmodesmata can respond to PAMPs. For this, we used the *choline transporter-like1* (*cher1*) mutant that is reported to have only simple plasmodesmata^24^. Microprojectile bombardment assays revealed that chitin triggered a significant reduction in the cell-to-cell mobility of GFP in mature leaves of *cher1* (Fig. S3D, Supplemental Dataset S2), suggesting that simple plasmodesmata can close in response to PAMPs.

To directly investigate the correlation between plasmodesmal structure and stress-induced closure we used Serial Block Face Scanning Electron Microscopy (SBF-SEM). We imaged the abaxial epidermal cell layer that we assay in microprojectile bombardment experiments and compared the abundance of simple and branched plasmodesmata between cells in young sink and young source leaves harvested 24 hours after covering of mature leaves (Fig. S3E, F, Movie S1). Comparison of the ratios of simple to branched plasmodesmata structures in young sink and young source leaves showed no significant difference between them (Fig. 3K). Therefore, we concluded that plasmodesmal responsiveness is not defined by plasmodesmal structure.

### Inducing plasmodesmal closure in young sink leaves enhances defence and reduces growth

Our finding that the carbon sink or source status of leaves defines whether plasmodesmata respond to stress suggests that young sink leaves prioritise phloem unloading over the enhancement of defence offered by plasmodesmal closure. If the loss of plasmodesmal closure is detrimental to overall defence, as observed in mature leaves of plasmodesmal mutants^2,3^, we expect that young leaves would be more susceptible than mature leaves to infection. To test this, we assayed for *Pseudomonas syringae pv. tomato* (*Pst*) DC3000 growth in young and mature leaves. Four days post inoculation, young leaves showed more signs of disease than mature leaves, with visible lesions and areas of chlorosis (Fig. 4A). Quantitation of bacterial growth identified significantly higher bacterial growth in young leaves than mature leaves (Fig. 4B). Therefore, young sink leaves are more susceptible to *Pst* DC3000 than source leaves, consistent with the possibility that they prioritise growth at the expense of defence.

**Figure 4:**
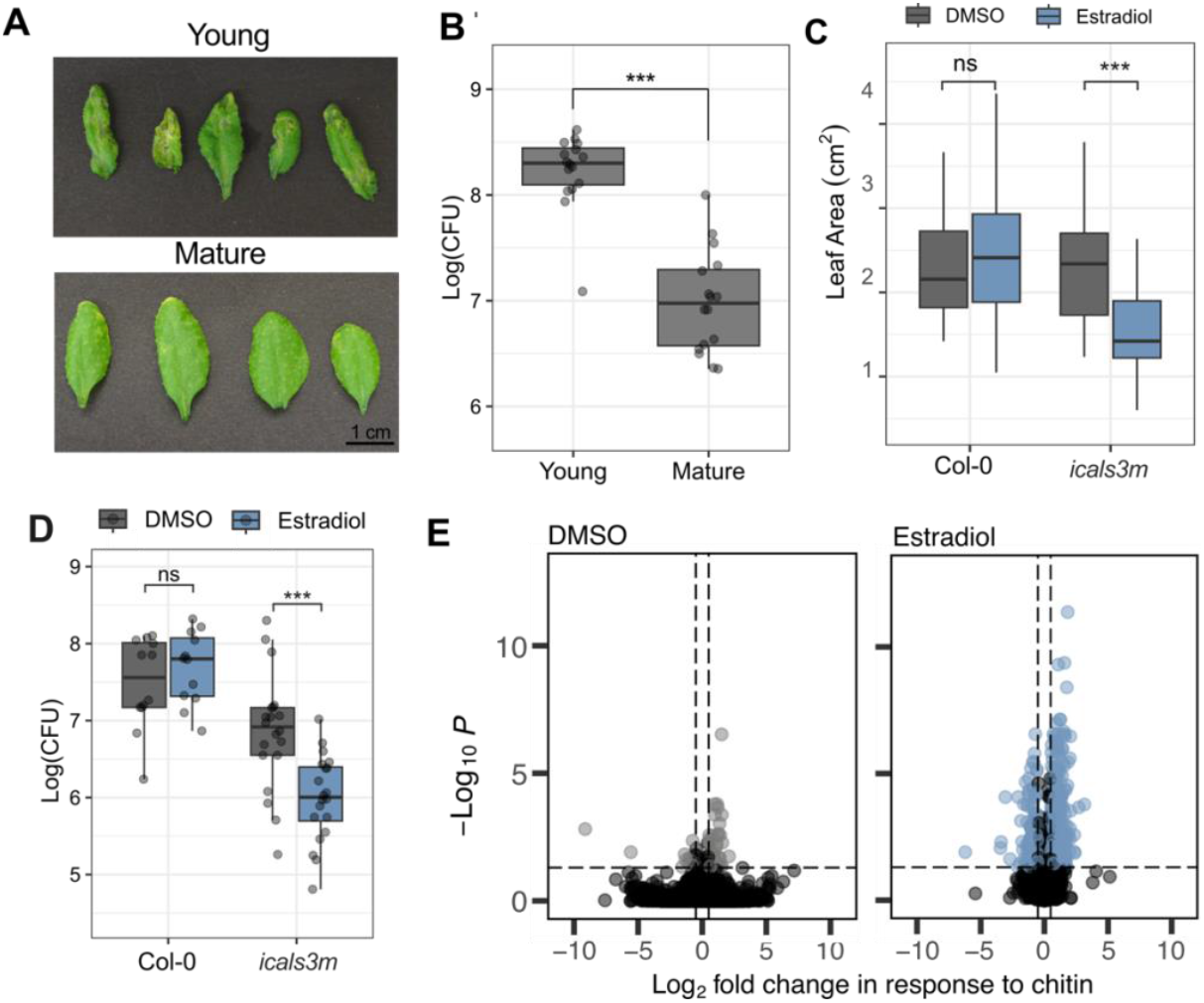
Plasmodesmal closure regulates both growth and defence Figure 4: Plasmodesmal closure regulates both growth and defence. (**A**) Young and mature leaves 4 days post-infection with *Pst* DC3000. (**B**) CFU from young and mature leaves 4 days post-infection with *Pst* DC3000, n = 16. (**C**) Quantification of the leaf area of young leaves of Col-0 and *LexA::icals3m* (icals3m) 4 days after estradiol or DMSO treatment, n = 12. (**D**) Colony Forming Units (CFU) from young leaves of Col-0 and *LexA:icals3m*, treated with DMSO (grey) or estradiol (blue), infected with *Pst* DC3000, points show individual replicates, n ≥ 12 (**E**) Volcano plots where blue and grey points indicate significantly differentially expressed genes in DMSO (left, grey) and estradiol (right, blue) treated *LexA::icals3m* respectively, where differential expression is defined by p_adj_<0.05 and log_2_(fold change) > |0.5| and black points indicate genes which are not significantly differentially expressed.. Asterisks indicate statistical significance compared to the mock treatment: *p < 0.05, **p < 0.01, and ***p < 0.001.

If this hypothesis is correct, we also expect that the induction of plasmodesmal closure in young sink leaves would both impair growth and enhance defence (and resistance). We previously generated a line in which we can induce plasmodesmal closure via estradiol-inducible production of callose with the over-active *icals3m* allele of CALLOSE SYNTHASE3^19,25,26^. We verified that estradiol treatment induced plasmodesmal closure in young sink leaves of *LexA::icals3m* (Fig. S4A,B, Supplemental Dataset S2) and subsequently assessed the effect of plasmodesmal closure on the growth of young leaves. Measuring leaf area four days after estradiol treatment revealed that induced leaves are smaller than uninduced leaves, indicating plasmodesmal closure in sink leaves impairs growth (Fig. 4C).

To explore how plasmodesmal closure affects defence and overall resistance to infection we inoculated young leaves of *LexA::icals3m* with *Pst* DC3000 24 hours after inducing plasmodesmal closure. Four days post-inoculation, estradiol-treated *LexA::icals3m* young leaves showed significantly less bacterial growth than DMSO-treated young leaves (Fig. 4D), suggesting that inducing plasmodesmal closure in *LexA::icals3m* enhances defence against *Pst* DC3000 in young leaves. Thus, growth and defence are negatively and positively correlated with plasmodesmal closure respectively in young leaves, consistent with the hypothesis that young leaves prioritise growth by not closing their plasmodesmata during immune responses.

Our experiments have suggested that the magnitude of the transcriptional response to chitin is correlated with plasmodesmal closure. Thus, we assessed transcriptional changes to chitin 24 hours after the induction of plasmodesmal closure in young leaves of *LexA::icals3m*. We identified genes that are differentially expressed in response to chitin when plasmodesmata are closed (*LexA::icals3m* estradiol treatment, chitin vs water) and when plasmodesmata are open (*LexA::icals3m* DMSO treatment, chitin vs water) and observed that there was a stronger transcriptional response to chitin treatment in young leaves when plasmodesmata are closed (503 genes upregulated, 298 genes downregulated) compared to when plasmodesmata are open (47 genes upregulated, 18 genes downregulated) (Fig. 4E and S4C, Supplemental Dataset S1). Again, while the number of chitin-induced DEGs was greater when plasmodesmata are closed, defence-related GO terms are enriched when both plasmodesmata are closed or open (Fig. S4C), suggesting it is the magnitude and not the nature of the response that is primarily altered.

## Discussion

Here, we observed that stress and immune responses are differentially executed in different tissues of the same plant. Across an Arabidopsis rosette, leaves that are still expanding execute a reduced response to PAMPs compared to mature expanded leaves. These differences in response are underpinned by the carbon status of the leaf - we observed that when young leaves are forced to transition from carbon sink to source they gain the ability to close their plasmodesmata, increase their transcriptional response to an immune elicitor and show increased resistance to a bacterial pathogen. Thus, our data implicate carbon metabolism as a regulator of the execution of immune responses (Fig. 4).

The physiological shift associated with the sink-to-source transition is from net consumers to net producers of carbon, changing the direction of carbon flow into and within the phloem. We reason that the sink-to-source transition may underpin the execution of immune-triggered plasmodesmal closure because young leaves prioritise phloem unloading above immune responses. This might represent a novel and specific trade-off between growth and defence; leaves that rely on carbon delivery via the phloem do not close their plasmodesmata despite the enhancement this offers to defence. Critically, comparing *Pst* DC3000 infection of young sink leaves with source leaves revealed that sink leaves are more susceptible, and execute a reduced transcriptional response to a PAMP, consistent with the idea that these leaves execute a compromised defence response.

We speculated that primary metabolism in young leaves might limit plasmodesmal response by reduced UDP-glucose availability, the substrate of callose synthases that ultimately mediate plasmodesmal closure. However, young leaves can produce macroscopic callose deposits in response to flg22 (Fig. 1), suggesting that substrate availability does not globally limit callose synthesis. Furthermore, the plasmodesmata between root cells (sink tissue) close in response to stress elicitors^27,28^, raising the possibility that the sink-source dependency of plasmodesmal responses might not hold in all plant organs or that externally supplied sucrose (commonly used for in vitro root growth) might alter immune responses.

Leaves of different ages have been demonstrated to differentially interact with defence hormone signalling^13,29^. These observations have lead to the proposal that, according to optimal defence theory, young leaves have higher value to plant survival^13^. While our findings here challenge this premise, these previous data led to the proposal that sugars might act as a signal that mediates tissue-specific response prioritisation^30^. Here, we have confirmed this possibility and identified carbon status and/or availability as a factor that defines the scale of immune responses executed by different tissues and overall defence success.

Our study has identified that the execution of some immune responses is dependent on the carbon metabolism status of a tissue. Both plasmodesmal responses and the magnitude of transcriptional responses in leaves are correlated with whether a tissue is a carbon sink or source, establishing primary carbon metabolism as a key regulator of the plant immune system. Whether this rests on the supply of carbon resources required to generate responses, or another regulatory mechanism, remains to be determined. Overall, our data further suggests that plasmodesmata balance the outputs of simultaneous cellular processes to optimise overall physiological success, leading to young leaves executing compromised defence while maintaining growth.

## Materials and Methods

### Plant materials and growth

The Columbia (Col-0) ecotype background of Arabidopsis was used as wildtype in this study. *LexA::icals3m*^19^, *p35s::MP17-GFP*^5^ and *cher1*^24^lines were in the Col-0 ecotype. All Arabidopsis seed was stratified for a minimum of 48 hours in the dark at 4°C before being sown onto John Innes F2 starter soil (100% Peat, 4kg/m^3^ Dolomitic limestone, 1.2Kg/m^3^ Osmocote Start). Soil-grown plants for experimental assays were grown in growth cabinets (Panasonic Versatile Environmental Test Chamber MLR-352-PE) with a short-day 10-hour light and 14-hour dark cycle. Experiments were performed on 5-week-old plants.

### Inducing transgene expression with estradiol in leaves

Transgene expression in *LexA::icals3m* was induced using estradiol treatment. To prepare the estradiol treatment, β-estradiol 17-acetate (estradiol, Sigma-Aldrich) was first dissolved in dimethyl sulfoxide (DMSO) and then diluted with distilled water containing 0.0001% Silwet L-77 (De Sangosse) to achieve a final concentration of 20 µM estradiol. A corresponding mock solution was prepared by diluting DMSO to a final concentration of 0.001% in water with 0.0001% Silwet L-77. Both abaxial and adaxial surfaces of young and mature leaves were painted with either the 20 µM estradiol solution or the 0.001% DMSO mock solution. Following treatment, plants were incubated for 24 hours prior to performing further assays.

### Microprojectile bombardment assays

Microprojectile bombardments were carried out as described^31^ where young and mature leaves were bombarded with 1 nm gold particles (BioRad) coated in pB7WG2.0-GFP and pB7WG2.0-RFP_ER_ using a Bio-Rad Biolistic PDS-1000/He particle delivery system (Bio-Rad, 1652257). Two hours post-bombardment, leaves were syringe-infiltrated with an elicitor treatment (dH_2_O, 500 µg/ml chitin, 100 nM flg22, 50 µM Abscisic acid, 100 µM Salicylic acid, or 10 mM hydrogen peroxide). For microprojectile bombardments involving estradiol treatment, leaves were treated with a 20 µM estradiol solution or a 0.001% DMSO mock solution 24 hours prior to performing bombardments. GFP movement was visualised 16-24 hours later using a Zeiss LSM800 confocal microscope with a 20× objective (Apochromat 20×/0.8 water immersion). GFP was excited at 488 nm using an argon laser and collected at 500-546 nm and RFP was excited at 561 nm using a DPSS laser and collected at 562-617 nm. After image acquisition, cell-to-cell movement of GFP was quantified. The number of nuclei showing GFP fluorescence was counted, where a count of one nucleus indicates GFP has not spread from the transformed cell. Where specified, GFP movement was shown normalised to the mean of mock treatments within each tissue age. Bombardment data was analysed using a median bootstrap approach to account for the non-normal distribution and unequal variance of data and p-values were corrected for multiple comparisons using Holm-bonferonni^32^.

### ROS assays

Leaf discs were harvested from mature and young leaves using a 5mm leaf corer. Six hours after harvesting a working solution (20 mg/ml peroxidase, 20 mM L-012, and d) was added to the leaf discs. Immediately before measurement, leaf discs were treated with either sterile distilled water, 100 nM flg22, or 500 ng/mL chitin. Emitted luminescence was measured for 90 minutes after the addition of elicitors by a Varioskan Flash Plate Reader (Thermo Fisher Scientific). The intensity of the first ROS peak was quantified by integrating the relative light unit (RLU) in the first 45 minutes of luminescence measurements to calculate the cumulative RLU. A total of twelve plants were sampled across three experimental replicates for each leaf age and treatment. From each plant, three leaf discs were harvested and the cumulative RLU for each plant was determined by taking the mean cumulative RLU of three leaf discs. Cumulative RLU was compared between leaf ages and treatments in a mean bootstrap analysis incorporating a multiple comparisons test ^32^.

### Macroscopic callose quantification

Young and mature leaves were syringe-infiltrated with either a mock (dH_2_O) or 100 nM flg22 solution. 22 to 30 hours post-treatment leaves were syringe infiltrated with 1% aniline blue in PBS buffer (pH 7.4). Aniline blue staining of callose deposition, was imaged using a Zeiss LSM800 confocal microscope with a 10× objective 10× objective (EC Plan-Neofluar 10×/0.30 M27-Air). Aniline blue was excited using a 405 nm argon laser and collected at 410-500 nm. Z-stacks from five sites were acquired per leaf, and maximum projections of these z-stacks were used in image quantification. Aniline blue stained macroscopic callose deposits were quantified using an automated image analysis pipeline^33^. For experiments comparing 100 nM of flg22 to water treatment, at least 16 leaves were imaged per leaf age and treatment combination across four experimental replicates. The data was analysed using a median bootstrap approach to account for the non-normal distribution and unequal variance of data with comparisons carried out within each leaf age ^32^.

### Phloem unloading of GFP in leaves

Changes in GFP unloading in young leaves were quantified using pAtSUC2::GFP lines expressing GFP under the promoter of SUC2, a sucrose H+ symporter specifically expressed in phloem companion cells^17^.Five-week-old *pAtSUC2::GFP* plants were covered in foil or left uncovered for either 24-30 or 48-52 hours. Young and mature leaves from uncovered plants were imaged alongside young leaves from covered plants. Both the left and right abaxial side of each leaf were imaged. Images of GFP unloading in young leaves from covered and uncovered plants were acquired using an Axio ZoomV16 stereomicroscope with a Zeiss Plan Z1.5×/0.25, monochrome camera ORCA FLASH4 (C13440, Hamamatsu), and SPECTRA Light Engine (Lumencor). Leaves were illuminated with 463 nm to 487 nm light and fluorescence was collected with a 525/50 nm band-pass filter. Z-stack projections were generated using the Zeiss Blue v2.6 built-in software for the extended depth of focus wavelets method. Images of GFP unloading were analysed using FIJI and R scripts described in Bellandi *et al*. ^17^ originally used to quantify CFDA unloading. The images taken on either side of a leaf were considered technical replicates and averaged, resulting in one mean peak width value per leaf. Each leaf was considered a single biological replicate. A minimum of 13 mature leaves and 20 young leaves were quantified for each tissue type and time point. A Wilcoxon rank sum test was used to compare the mean widths of peaks for each leaf between different tissue types. p-values were adjusted for multiple comparisons using the Holm correction in the ggpubr R package (Holm^34^; v0.0.6^35^).

### Quantification of starch and soluble sugars

Young and mature leaves were harvested from covered and uncovered plants at the end of the day, either 24 or 48 hours after wrapping treatment. Leaf tissues were harvested at the end of day. Immediately after harvesting, leaves were weighed and flash-frozen in liquid nitrogen. Each young sample consisted of six harvested young leaves pooled from three different plants. Each mature sample consisted of three mature leaves pooled from three separate plants. Each sample was considered a biological replicate. For the 24-hour timepoint, a total of 18 samples were pooled across three experimental replicates for each tissue age and wrapping treatment. For the 48-hour timepoint, a single experimental replicate was carried out with 6 samples per tissue type. To extract starch and soluble sugars, leaf material was first homogenised in 0.7M perchloric acid using a mix mill (Retsch). Homogenised samples were pelleted by centrifugation with the insoluble pellet used to quantify starch as outlined^36^. The supernatant was used to quantify soluble sugars by first neutralising samples to a pH of 6-7 using neutralisation buffer (2 M KOH, 400 mM MES) and quantifying sugar content in a solution of 1M HEPES.KOH (pH 7.6), 1 mM ATP, 1 mM NAD, 100 mM MgCl_2_, 1.4U Hexokinase and H_2_O. An Omega FLUOstar microplate reader was used to quantify soluble sugars in an enzymatic assay. Glucose, fructose, and sucrose contents were determined by measuring sample absorbance after adding glucose-6-phosphate dehydrogenase, phosphoglucose isomerase and invertase, respectively. The mean absorbance of three technical replicates was used to calculate soluble sugar contents in mg per g of fresh weight. Soluble sugar and starch contents were then analysed using an ANOVA, with significant differences between individual tissue types determined in a post-hoc Tukey HSD test.

### Leaf area measurements

The four youngest leaves of five-week-old Col-0 and *LexA::icals3m* plants were painted abaxially and adaxially with 20 µM estradiol or 0.001% DMSO (mock). Four days post-treatment, leaf area was measured. The curling of leaves after estradiol treatment made measuring leaf area whilst attached challenging, and so leaves were detached and flattened, so the whole leaf area could be photographed. Leaf area was quantified from images using FIJI^37^. A total of four plants were quantified per genotype, with four young leaves measured per plant. A linear mixed model was conducted to determine the effect of treatment (estradiol vs DMSO), genotype (Col-0 vs *LexA::icals3m*) on leaf growth with leaf number included as a nested variable. Each leaf was considered a replicate. Following the three-way ANOVA, a Tukey HSD test was carried out to determine significant differences between treatments, genotypes, and leaf numbers.

### Quantification of PSII efficiency

The operating efficiency of PSII (ΦPSII) can be calculated using chlorophyll fluorescence emission. As defined by Genty et al. ^38^, ΦPSII can be calculated as Φ(PSII) = (Fm′−F′)/Fm′ where F is defined as steady-state chlorophyll fluorescence under actinic light conditions and Fm’ is the maximum chlorophyll fluorescence emission in light-adapted tissues when reaction centres are closed. F’ can be determined from chlorophyll fluorescence in actinic light conditions, whilst Fm’ can be determined by exposing light-adapted leaves to a saturating light pulse. To determine ΦPSII in the growth conditions used here, chlorophyll fluorescence was measured using an IMAGING-PAM Maxi Chlorophyll Fluorometer (Heinz Walz GmbH). First, Col-0 plants were dark-adapted for a minimum of 45 minutes, and then young and mature leaves were detached and placed in the IMAGING-PAM. Leaves were first exposed to a saturating light pulse of 845 µmol m^-2^s^-1^ and then actinic light was set to match growth cabinet conditions at 189 µmol m^-2^s^-1^. Every 60 seconds a saturating light pulse was used to measure Fm’. After approximately 20 mins of acclimation to actinic light, the value of Fm’ reaches its maximum, at which point Fm’ and F’ can be used to calculate ΦPSII under the actinic light of 189 µmol m^-2^s^-1^. ΦPSII was quantified in both young and mature uncovered and covered leaves and from 5-week-old Col-0 plants. Chlorophyll fluorescence was measured in young sink and young source leaves 24 hours after covering mature leaves with foil. In total, twenty-eight young leaves and twelve mature leaves were measured across two experimental replicates. A linear mixed model analysis to compare young sink and young source leaves was performed with experimental replicate included as a nested variable using the lmerTest R package^39^.

### RNA extraction and sequencing

To compare transcriptional response to chitin across tissue age, mature leaves from five-week-old Col-0 lines were syringe infiltrated with mock (water and 0.00001% Silwet L-77) or chitin (500 µg/ml chitin, water and 0.00001% Silwet L-77) and harvested 30 minutes later. Three mature leaves were harvested from three plants pooled together into one mature leaf sample. Young leaves were defined as leaves two and three and six young leaves from three plants were pooled into one young leaf sample. Each pooled sample was considered a single biological replicate, with three biological replicates sampled per leaf age and elicitor combination and sequenced by Novogene.

The transcriptional response to inducing plasmodesmal closure and chitin treatment was compared in Col-0 and *LexA::icals3m* lines. Young leaves of Col-0 and *LexA::icals3m* lines were painted with 20 µM estradiol or 0.001% DMSO (mock). 24 hours later leaves were syringe-infiltrated with a chitin (500 µg/mL chitin in distilled water) or mock (distilled water) solution and sequenced by Genewiz (Azenta life sciences). Leaf tissue samples were harvested 1 hour after elicitor treatment. Six young leaves from three plants were pooled per biological replicate, and five biological replicates were collected for each treatment-elicitor-genotype combination.

The transcriptional response to chitin treatment was compared in young sink and young source. Mature leaves of 5-week-old Arabidopsis Col-0 were covered or left uncovered. 24 hours later young leaves from covered and uncovered plants were syringe-infiltrated with a chitin (500 µg/mL chitin in distilled water) or mock (distilled water) solution. Leaf tissue samples were harvested 1 hour after elicitor treatment. Six young leaves from three plants were pooled per biological replicate, and five biological replicates were collected for each treatment-elicitor-carbon status combination. RNA samples were sequenced by Genewiz (Azenta life sciences).

### Gene expression analysis

Reads for all datasets were checked for quality using FastQC (v0.11.8^40^). Adaptors were trimmed and low quality reads with a phred of less than 20 were removed using TrimGalore (v.0.5.0^41^) and Cutadapt (v.1.7^42^). Reads were then mapped against the TAIR10 reference Arabidopsis genome using Hisat2 (v.2.1.0^43^). SAMtools was used to convert files for input into StringTie, which was used calculate read counts and create a matrix of counts^44,45^. Raw read counts were used in differential gene expression analysis using DESeq2 (v.1.38.3^46^) in R studio (v.4.3.3). Wald tests were performed on transcriptional data to extract differences in gene expression between elicitors (mock vs chitin), treatments (DMSO vs estradiol, where applicable), and/or genotypes (Col-0 vs icals3m, where applicable). Differentially expressed genes were identified from the Wald test based on a log_2_[fold change] of greater than |0.5| and an adjusted p-value of less than 0.05. Additional R packages were used to carry out data analysis: GO-term enrichment analysis was carried out using TopGO (v.2.54.0^47^) and enrichment score was calculated by taking the −log_10_ of the calculated fisher value; Heatmaps were built using ComplexHeatmap (v.2.18.0^48^); Volcano plots were plotted using EnhancedVolcano (v1.20.0^49^); shared differentially expressed genes were identified using RVenn (v.1.1.0^50^).

### Imaging of Citrine-LYM2 localisation

Young leaves of *lym2*.*LYM2pro::Citrine-LYM2* plants were detached from 5-week-old plants and immediately imaged to confirm the expression of *Citrine-LYM2* in young leaves^3^. Images were acquired using a Zeiss 800 confocal microscope with a 63× objective (C-Apochromat 63x/1.20 W Korr UV VIS IR). LYM2-Citrine was excited using a 488 nm laser and collected at 500-617 nm. Citrine-LYM2 expression was screened across more than 10 young leaves on three separate days to determine there was no leaf-to-leaf variation in localisation.

### Pathoassay with *Pseudomonas syringae* DC3000

Pathoassays with *Pseudomonas syringae* pv. *tomato* (*Pst*) DC3000 were carried out by syringe infiltrating young and mature leaves with a 5 ×10^4^ CFU/mL *Pst* DC3000 suspension. Four days post-inoculation leaf discs were harvested using a 5 mm corer. Six leaf discs from three plants were pooled during harvesting to make up one biological replicate. The experiment was repeated three separate times. Leaf discs were lysed using a Geno/Grinder (SPEX SamplePrep2010) and a dilution series was carried out to determine bacterial growth.

For experiments investigating the impact of inducing plasmodesmal closure on *LexA::icals3m* the three youngest leaves (leaves one to three) of *LexA::icals3m* and Col-0 plants were painted with 20 µM estradiol or 0.001% DMSO 24 hours before inoculation. Data was analysed using a linear mixed-effects model comparing the impact of genotype and treatment on the log of CFU, with the experimental replicate included as a nested random variable using the lmerTest R package^39^. Significant differences between genotypes and treatments were determined in a post-hoc Tukey pairs comparison.

In the assay comparing Col-0 young and mature leaf susceptibility, the experiment was replicated three times, with four biological replicate samples per experiment for a total of twelve samples. Data was analysed using a linear mixed-effects model comparing the impact of treatment on the log of CFU, with the experimental replicate included as a nested random variable using the lmerTest R package^39^.

### Serial block face-scanning electron microscopy

Young leaves from covered and uncovered plants were fixed in 1.5% (v/v) glutaraldehyde, 2.5% (v/v) paraformaldehyde, 5% (v/v) sucrose, 0.01% (v/v) IGEPAL in 0.1 M sodium cacodylate buffer (pH 7.4). Fixation and subsequent processing were carried out were inside a Pelco BioWave Pro+ (Agar Scientific) under vacuum. After fixation, samples were rinsed in 0.1M sodium cacodylate buffer before post-fixation in 2% osmium tetroxide in 0.1M sodium cacodylate buffer. Samples were then treated with 2.5% potassium ferrocyanide in 0.1M sodium cacodylate buffer, rinsed with water, incubated in 1% unbuffered thiocarbohydrazide, rinsed again with water, and finally immersed in 2% aqueous osmium tetroxide, followed by a final water rinse.

Following heavy metal staining, samples were dehydrated through a graded ethanol series (20%, 50%, 75%, and 3 x 100%) and infiltrated with Durcupan ACM resin using a graded series of concentrations in ethanol (25%, 50%, 75%, 90% and 3 x 100%). Durcupan ACM resin was prepared from epoxy resin (22.8g), hardener 964 (20g), accelerator 960 (0.6g) and dibutyl phthalate (0.2g). After the final infiltration, samples were transferred to 100% fresh resin, rotated overnight in a fume hood, and embedded in 13mm flat TAAB cavity moulds with fresh resin, which was polymerised at 60°C for 48h.

Samples were manually trimmed to size using a razor blade and then adhered to a Gatan 3View specimen pin, standard type 1.4 mm flat (Agar Scientific), using conductive epoxy glue. A Leica ARTOS 3D Ultramicrotome (Leica microsystems) was used with a DiATOME trim 45 diamond trimming tool (Leica microsystems) to refine the geometry of the resin block. Sample pins were subsequently sputter coated with 15 nm of platinum using a Leica ACE 600 sputter coater (Leica microsystems). The samples were placed in a Zeiss Gemini 300 SEM (Carl Zeiss Microscopy) with focal charge compensation, integrated with a Gatan 3View® 2XP in situ ultramicrotome and an OnPoint detector (AMETEK). Gatan DigitalMicrograph DMS3 software (AMETEK) was used to capture images in high vacuum conditions at 1.5kV. Sections were taken at a thickness of 50 nm, with a pixel size of 30 nm, pixel dwell time of 8 µs, and image size of 2048 × 2048. Datasets were aligned and segmented using Microscopy Image Browser (MIB) (https://mib.helsinki.fi, Microscopy Image Browser), and visualized using Amira software (Thermo Fisher Scientific). Plasmodesmata were classified as simple when they had only one opening on either side of the cell wall and branched when they had more than one opening on either side of the cell wall.

## Acknowledgements

We acknowledge access to the John Innes Centre Bioimaging Facility. We acknowledge the Seung lab for technical assistance in the sugar and starch measurements and David Seung for critical conversations and advice throughout the project. We would like to thank Núria Real-Tortosa for critical reading of the manuscript.

Work in the Faulkner lab has been funded by the Biotechnology and Biological Science Research Council (Grants: BB/X010996/1; BB/X007685/1; BB/Y008782/1; BB/X016056/1) and the European Research Council (Grant: 725459 - “INTERCELLAR”). ER was supported by a John Innes Foundation Rotation PhD studentship.

## Supplemental Data

Supplemental Figures 1-4

Supplemental Supplemental Dataset S1: Lists of differentially expressed genes in all experiments in this study

Supplemental Supplemental Dataset S2: Lists of cell counts (no. cells showing GFP) for all microprojectile bombardment experiments in this study

Supplemental Movie 1: Serial SEM sections from a young sink leaf showing annotation of plasmodesmata (purple) and the cell wall (yellow).

