## Supplemental Figures for "Primary metabolism underpins the execution of immune responses in different tissues of the same plant"

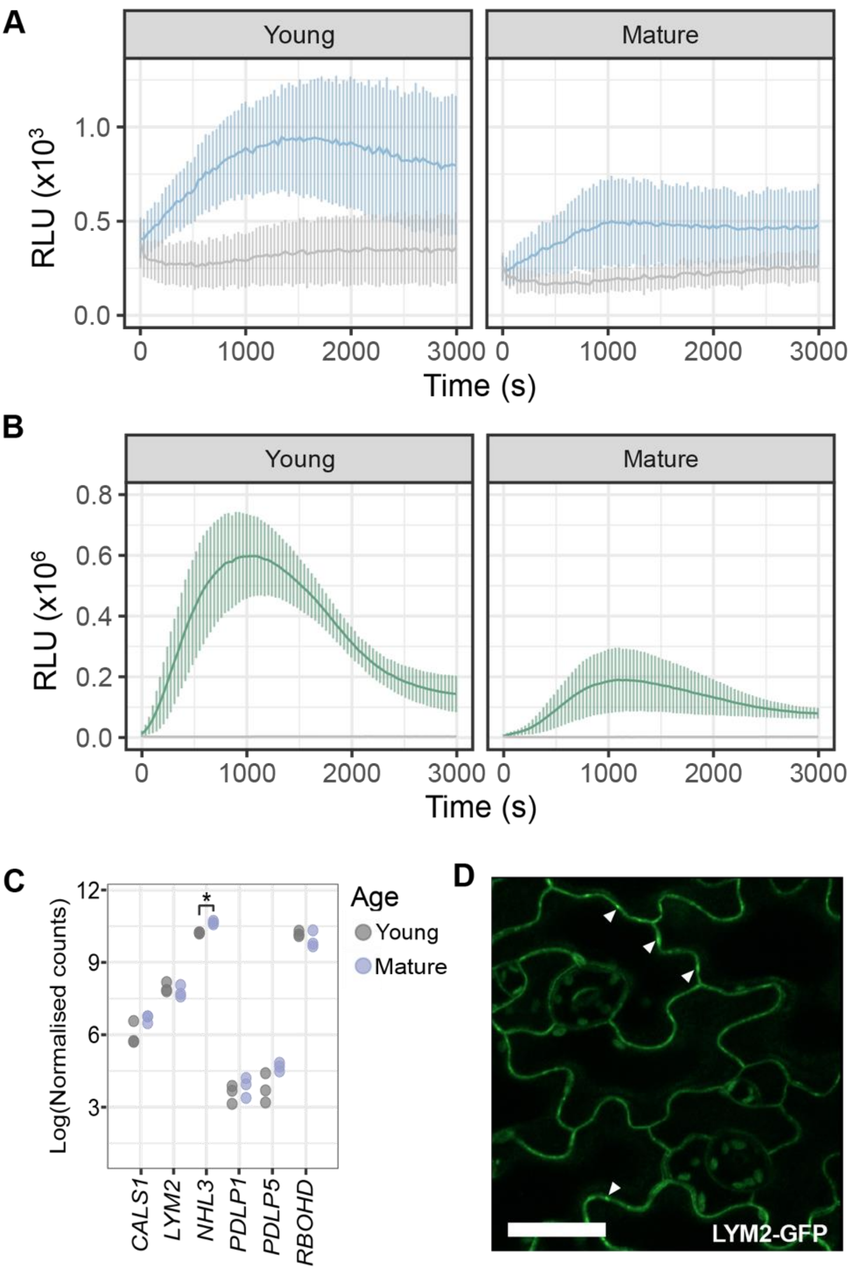


**Figure S1: Differential expression of plasmodesmal-related signaling components cannot explain differences in plasmodesmal regulation**. – (**A**) Relative light units (RLU) of leaf discs from young and mature leaves in response to mock (grey) versus chitin (blue) over time and (**B**) in response to mock (grey) versus flg22 (green) over time. (**C**) Normalised gene counts of plasmodesmal signalling-related genes expressed in young and mature samples in mock-treated samples, where "*" indicates an adjusted p-value < 0.05 for significantly differentially expressed genes between young and mature mock-treated tissue. (**D**) *pAtLYM2::LYM2-citrine* expression in young leaves where the arrows indicate the localisation of LYM2-GFP to plasmodesmata (n = 10), the scale bar indicates 200 μm.

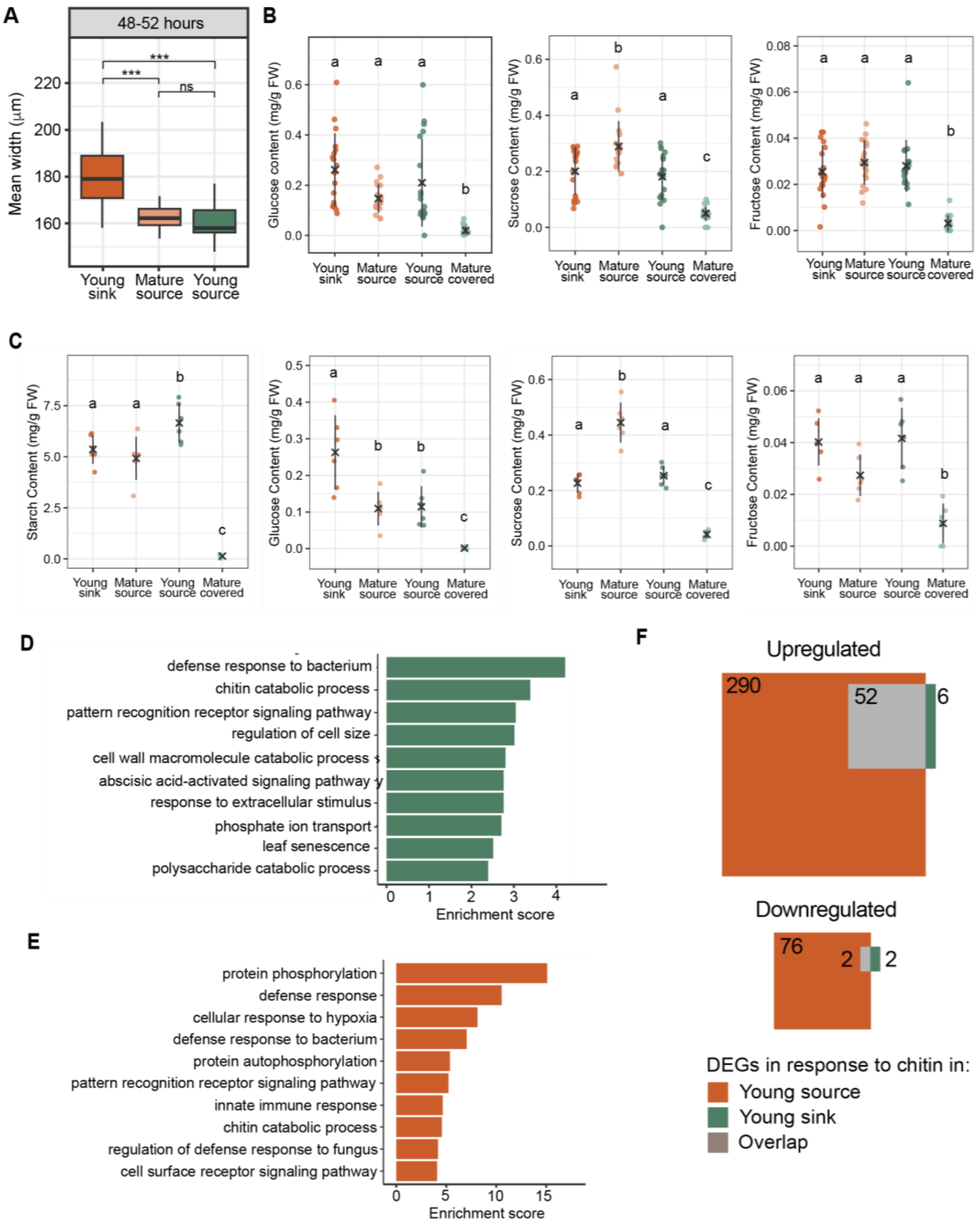


**Figure S2: Young source leaves show physiological and gene expression differences compared to young sink leaves** – (**A**) Mean width of GFP unloading in young sink (dark orange, n ≥ 20), mature source (light orange, n ≥ 13), and young source (dark green, n ≥ 20) leaves 48-52 hours after covering or leaving uncovered in *pAtSUC2::GFP* lines. “ns” indicates no significant difference and ’***” indicates $p < 0.001$. (**B**) Glucose, sucrose, and fructose content measured as mg/g of fresh weight (FW) 24 hours after foil wrapping mature laves of covered plants (n = 18). (**C**) Starch, glucose, sucrose, and fructose content 48 hours after foil wrapping mature leaves of covered plants (n = 6). Different letters indicate significant differences between tissue types where $p < 0.05$, crosses indicate the mean value, and lines indicate +/- standard deviation. Points represent individual samples. Enriched GO-terms for differentially expressed genes upregulated in response to chitin in (**D**) young sink and in (**E**) young source tissue. (**F**) Venn diagram showing overlap between differentially expressed genes in young sink and young source tissue (log2[fold change] > |0.5|, padj< 0.05).

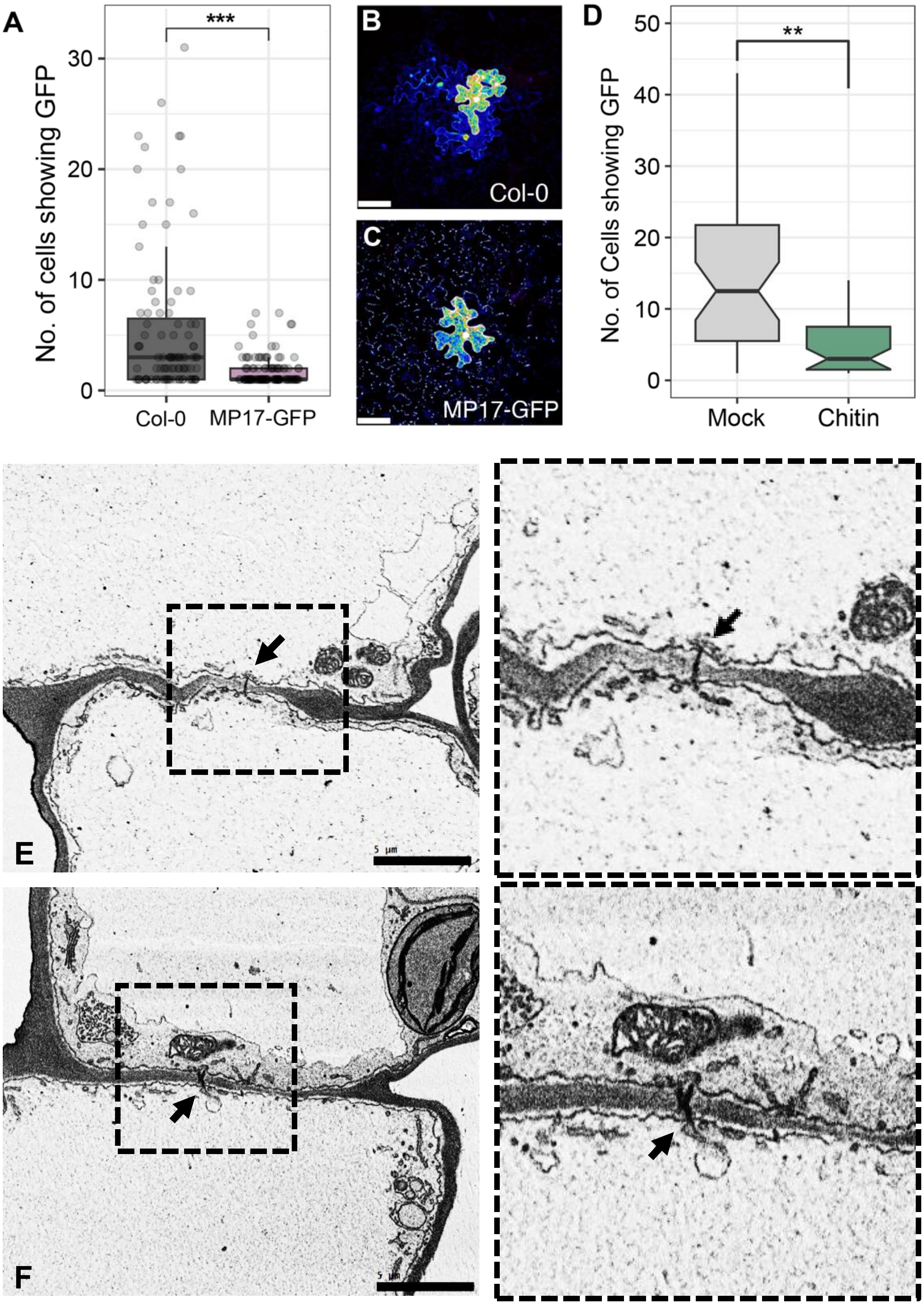


**Figure S3: Plasmodesmal structure does not determine plasmodesmal PAMP response** – (**A**) Cell to cell movement of GFP in microprojectile bombardment assays on Arabidopsis lines constitutively expressing MP17-GFP (n > 99). Confocal images of bombardment sites in (**B**) Col-0 and (**C**) MP17-GFP showing localisation of MP17-GFP at plasmodesmata and cell-to-cell movement of free GFP. Scale bars indicate 50 µm. (**D**) Boxplots of microprojectile bombardments in *cher1* lines after mock (water) or chitin treatment, n > 36. "**" indicates p < 0.01. (**E-F**) SBF-SEM image of a (**E**) simple and (**F**) branched plasmodesma with the dotted boxes enlarged on the right. Plasmodesma are indicated by black arrows. Scale bars are 5µM.

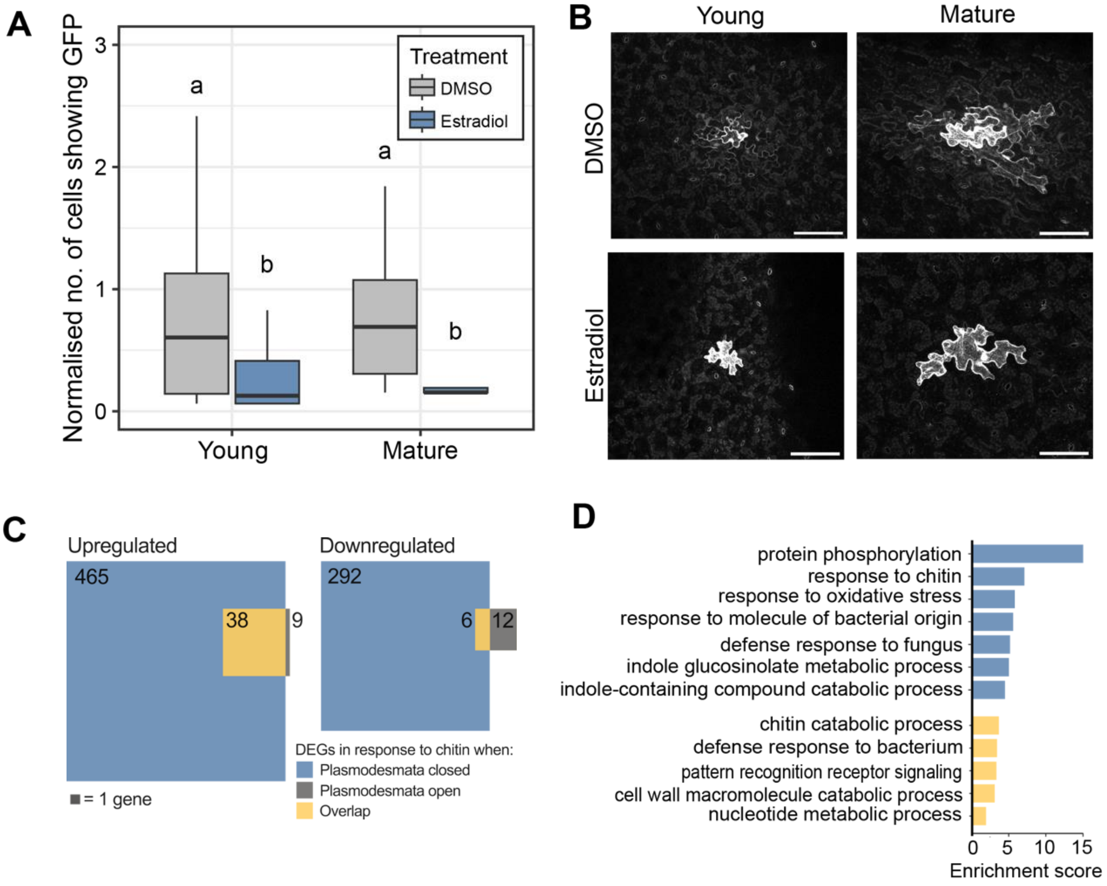


**Figure S4: Inducing plasmodesmal closure in young leaves of *LexA::icals3m* lines alters the magnitude of the transcriptional response to chitin** – (**A**) Cell-to-cell movement of GFP in *LexA::icals3m* leaves treated with either DMSO (grey) or estradiol (blue). Different letters indicate significant differences in a bootstrapping analysis of the medians, $p < 0.05$, $n > 39$. (**B**) Confocal z-stack projections of GFP movement in DMSO- and estradiol-treated young and mature leaves. Scale bars indicate 100 µm. (**C**) Venn diagrams showing the numbers of chitin-triggered differentially expressed genes (DEGs) genes in young leaves of *LexA::icals3m* when plasmodesmata are closed (estradiol, blue), when open (DMSO, grey), and genes differentially expressed in both conditions (yellow) (log2[fold change] > |0.5|, padj< 0.05). (**D**) Enriched GO-terms for significantly upregulated genes in response to chitin in *LexA::icals3m* when plasmodesmal closure is induced (blue), when plasmodesmata are open (grey), and genes differentially expressed in both sets (yellow) ($\log_2$ fold change > |0.5|, adjusted p-value < 0.05
